# Using genetics to differentiate patients with similar symptoms: application to inflammatory arthritis in the rheumatology outpatient clinic

**DOI:** 10.1101/813220

**Authors:** Rachel Knevel, Saskia le Cessie, Chikashi C. Terao, Kamil Slowikowski, Jing Cui, Tom W.J. Huizinga, Karen H. Costenbader, Katherine P. Liao, Elizabeth W. Karlson, Soumya Raychaudhuri

## Abstract

Slow developing complex diseases are a clinical diagnostic challenge. Since genetic information is increasingly available prior to a patient’s first visit to a clinic, it might improve diagnostic accuracy. We aimed to devise a method to convert genetic information into simple probabilities discriminating between multiple diagnoses in patients presenting with inflammatory arthritis.

We developed G-Prob, which calculates for each patient the genetic probability for each of multiple possible diseases. We tested this for inflammatory arthritis-causing diseases (rheumatoid arthritis, systemic lupus erythematosus, spondyloarthropathy, psoriatic arthritis and gout). After validating in simulated data, we tested G-Prob in biobank cohorts in which genetic data were linked to electronic medical records:

- 1,200 patients identified by ICD-codes within the eMERGE database (n= 52,623);
- 245 patients identified through ICD codes and review of medical records within the Partners Biobank (n=12,604);
- 243 patients selected prospectively with final diagnoses by medical record review within the Partners Biobank (n=12,604).

The calibration of G-Prob with the disease status was high (with regression coefficients ranging from 0.90-1.08 (ideal would be 1.00) in all cohorts. G-Prob’s discriminative ability was high in all cohorts with pooled Area Under the Curve (AUC)=0.69 [95%CI 0.67-0.71], 0.81 [95%CI 0.76-0.84] and 0.84 [95%CI 0.81-0.86]. For all patients, at least one disease could be ruled out, and in 45% of patients a most likely diagnosis could be identified with an overall 64% positive predictive value. In 35% of instances the clinician’s initial diagnosis was incorrect. Initial clinical diagnosis explained 39% of the variance in final disease prediction which improved to 51% (*P*<0.0001) by adding G-Prob genetic data.

In conclusion, by converting genotypes into an interpretable probability value for five different inflammatory arthritides, we can better discriminate and diagnose rheumatic diseases. Genotypes available prior to a clinical visit could be considered part of patients’ medical history and potentially used to improve precision and diagnostic efficiency in clinical practice.

## INTRODUCTION

The prevalence of patients with whole-genome genotyping data is dramatically increasing (1–3); genome-wide genetic data are being collected as part of biobanking efforts, routine care, and direct to consumer genotyping. Genotype data provides a patient-specific, time-independent risk profile. Once available, genetic data could be interpreted in the context of signs and symptoms to prioritize different diagnoses. Without signs or symptoms, these data may not be particularly informative for most complex rheumatic diseases since these diseases tend to be rare(4–11). But, when genetic data are available at an initial visit, they could be used in ongoing clinical care in real-time(12, 13). In this study, we explored whether genetic data can facilitate disease differentiation in patients with similar early-disease-stage symptoms of inflammatory arthritis at their first visit to the outpatient clinic.

We developed G-Prob (genetic-probability-tool), a tool that can facilitate discrimination between multiple diseases by providing probabilities of likely diseases based on patients’ genetic data (Fig. 1). The advantage of providing probabilities instead of a binary diagnostic classification is that it synergizes with the probabilistic reasoning of clinicians and can easily be incorporated into a differential diagnosis. In instances where the genetic probability for a disease is sufficiently high or low, a clinician can prioritize diagnostic testing accordingly.

**Fig. 1.**
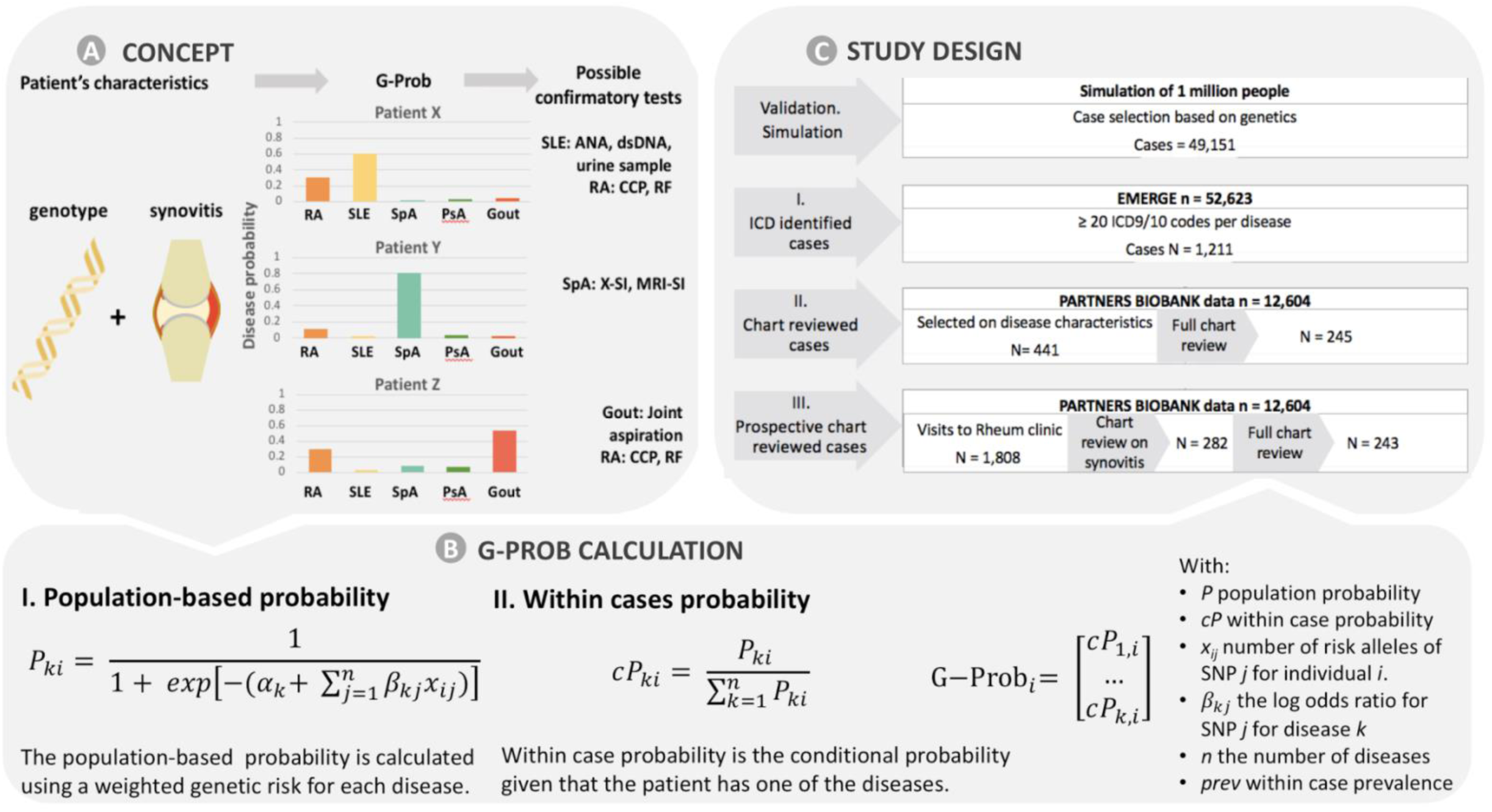
Schematic depiction of G-Prob and study design. Panel A schematically depicts the principle of G-Prob. In patients with symptoms (in this study: inflammatory arthritis which means a suggestion of synovitis at clinical exam in this example) the G-Prob gives a probability for each of the possible diseases. The magnitude and difference between disease probabilities can guide clinicians to subsequent tests. For instance, relevant tests for patient X would focus on differentiating SLE from RA such as autoantibody testing and urine examination, imaging could be prioritized in patient Y, a joint aspiration in patient Z would be relevant and the G-Prob algorithm suggests that no additional tests are needed to rule out SLE in patients Y and Z. B. The genetic probability calculation consists of two steps: first it measures the population-based probability using a weighted genetic risk score. Next, it obtains the within patients’ probabilities normalizing the population-based probabilities. The end product, G-Prob, gives each patient a probability for each disease. C. This study contained multiple validation and testing phases summarized by this flow-chart

We tested the performance of G-Prob in the setting of the rheumatology outpatient clinic, where many patients present with inflammatory arthritis (a typical swelling an tenderness of joints that rheumatologist identify at physical examination and that correlates with synovitis, which is inflammation of a membrane covering the joint (synovium)) as the first symptom of inflammatory arthritis. Approximately 80% of the patients with inflammatory arthritis in the outpatient rheumatology clinic are eventually diagnosed with rheumatoid arthritis (RA) (14, 15), systemic lupus erythematosus (SLE) (16), spondyloarthropathy (SpA) (17–19), psoriatic arthritis (PsA) (20) or gout (21). Since many risk loci have been identified for these diseases(22–29) and inflammatory arthritis is not present in healthy individuals, the population of people with inflammatory arthritis is ideal to test our hypothesis. Such patients are often misdiagnosed at their first visit. If the correct specific diagnosis for patients with inflammatory arthritis could be more quickly defined, it will be possible to start therapies more quickly. Many studies have clearly demonstrated that early diagnosis and treatment prevents disability and permanent damage in inflammatory arthritis (30–37). Equally importantly, the ability to define the correct disease early also prevents patients from being treated with inappropriate immunomodulatory therapies.

## RESULTS

G-Prob uses genetic information (combined in a genetic risk score (GRS)) from multiple diseases to calculate patients’ conditional probabilities for each disease, assuming that one of the diseases is present. In our proof-of principle study we utilized the G-Prob method to calculate the probabilities for RA, SLE, PsA, SpA and gout, using both single-nucleotide polymorphisms and HLA variants of uncorrelated risk single nucleotide polymorphisms (SNPs), as reported for European samples as well as the sex dependent population risk (Material and Methods).

First, we tested whether genetic information on its own is informative for disease discrimination. We simulated a population of 1 million individuals and assigned 49,151 cases with one of our five synovitis-causing diseases (Material and Methods and **Fig. S1**). We wanted to assigned individuals to have a disease or not, stochastically taking account the differences in their underlying genetic risk. Hence, we assigned case status when a sample’s genetic population-based disease risk (for one of the five diseases) was higher than a randomly sampled probability between zero and one. Since we assumed a disease prevalence of 1% for each disease in the simulated population, each disease was equally represented in our case set. This resulted in a simulated population of cases solely defined by genetics and a random factor, creating the ideal test setting for G-Prob.

G-Prob is a multiclass classifier, calculating probabilities for several diseases. For each disease, we can test if the probabilities match the true disease status. If G-Prob performs well, the patient’s true disease status should correspond with a high probability for that specific disease, while the probabilities for the other diseases should be low. We will further describe probabilities that concern a patient’s real disease as disease matching and probabilities that concern one of the other diseases as non-disease matching (see Materials and Methods).

Within the simulated cases, we observed that the distribution of the genetic probabilities matching the patients’ eventual diseases were clearly higher than the genetic probabilities for the other diseases (Fig. 2A, **Fig. S10**). Next, we tested whether the magnitude of the probabilities calibrated well with the real disease (based on application of the rheumatology classifications on complete follow-up). For this, we estimated the regression line between the probabilities and disease match (yes/no) using linear regression with the intercept constrained to zero, whereby the resulting beta (regression coefficient) gave the exact calibration (Material and Methods). Given that only genetic information influenced the case status, the beta was high in the simulated cases (beta=1.01, Fig. 2B). Whether G-Prob is useful to prioritize patients depends partly on how often the probabilities are highly informative values, that is low or high probabilities. In the simulated data, 32% of all probabilities were ≤0.05, making their corresponding diseases very unlikely. This cut-off corresponded to an NPV (negative predictive value) of 0.98 (Table 1) and may allow certain diagnoses to be effectively removed from the differential diagnosis in the right clinical context. At a patient level, 90% of the patients had at least one probability ≤0.05. On the other extreme, 64% of patients had a diagnosis with at least one probability ≥0.5; this corresponded to 13% of the assigned probabilities (PPV (positive predictive value) =0.70). In these instances, a diagnosis may be favored on the basis of genetics alone.

**Fig. 2.**
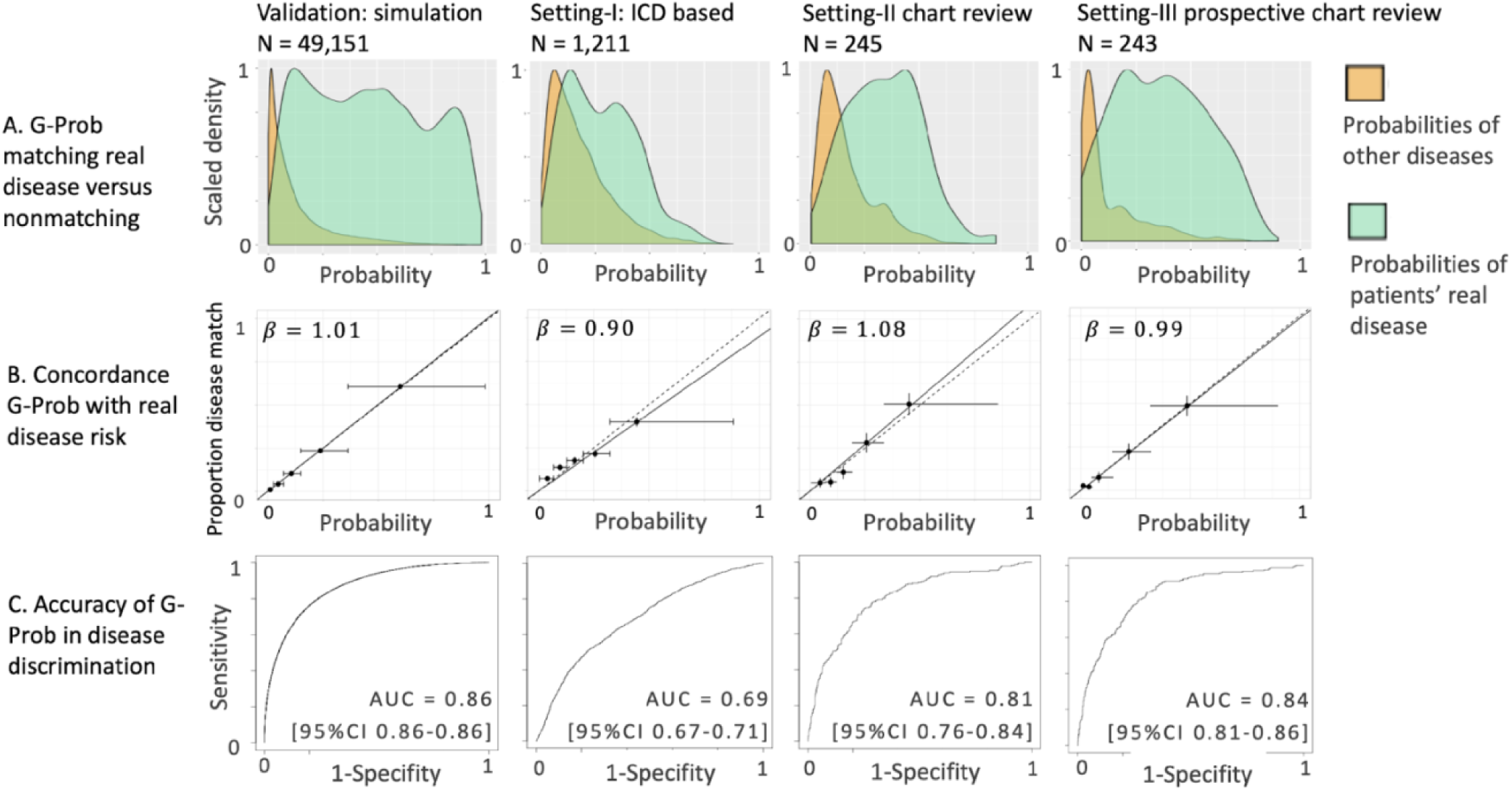
Discriminative ability of G-Prob and the concordance of predictions with observed disease occurrence. G-Prob gives each patient a probability for each disease that is part of the G-Prob. The performance of G-Prob is measured by comparing the magnitude of the probabilities with whether the probabilities match the real disease or one of the other diseases. In the simulation and setting I and II G-Prob contains five categories: rheumatoid arthritis, systemic lupus erythematosus, psoriatic arthritis, spondyloarthropathy and gout. In setting III, “Other” is added as sixth category. Row A. depicts the distribution of probabilities of the real diseases (green) and the probabilities of other diseases (orange). See **Fig. S10** for the depiction on the individual disease level. Row B. shows the concordance of G-Prob with patients’ real disease status. We pooled the disease probabilities and created a binary variable indicating whether the probability referred to the patient real disease or not. For the clarity of this graphs, we binned the probabilities of G-Prob into five equally sized bins. The x-axes display the mean G-Prob with range and the y-axis the proportion of the disease match with 95%CI confidence interval. The solid line and the *β* present the slope and 95%CI confidence interval of G-Prob with disease match as calculated with linear regression model. In the case of a perfect test performance, the solid line would lie exactly on the dashed diagonal line. Row C. depicts the receiver operating curve (ROC) which describes the balance between the sensitivity and the specificity and thereby the overall discriminative ability of G-Prob. The area under the curve (AUC) summarizes the area under the curve of the pooled data from all disease. See **Fig. S7** for the depiction on the individual disease level.

**Table 1.** Performance of G-Prob in ruling out and pointing towards diseases. Panel A shows the negative predictive value of the probabilities when we apply exclusion cut-offs of 0.05 and 0.2. A G-Prob below 0.05 is considered highly likely not to concern the probability of the patient’s real disease. Given that G-Prob constitutes of five diseases, 0.2 is the expected probability for each disease if genetics would not play a role in any of the diseases. Panel B gives the positive predictive value of the probabilities higher than 0.2 and 0.5. A probability higher than 0.5 is by definition the largest disease probability a patient has and would therefore strongly suggest the patient has that particular disease. Since each patient has multiple probabilities the table provides the percentage of probabilities as well as the percentage of patients that have probabilities above or below the cut-offs. More extensive test characteristics are depicted in **Fig. S11**.

| cut-off | G-Prob performance |  |  |  |  |  |  |  |  |  |  |  |
| --- | --- | --- | --- | --- | --- | --- | --- | --- | --- | --- | --- | --- |
|  | A. ruling out diseases |  |  |  |  |  | B. pointing towards diseases |  |  |  |  |  |
|  | ≤0.05 |  |  | ≤0.2 |  |  | ≥0.2 |  |  | ≥0.5 |  |  |
|  | % of pts | % of probs | NPV | % of pts | % of probs | NPV | % of pts | % of probs | PPV | % of pts | % of probs | PPV |
| **Simulation: Validation** | 90% | 32% | 0.98 | 100% | 67% | 0.93 | 100% | 33% | 0.47 | 64% | 13% | 0.70 |
| **Setting-I** | 51% | 13% | 0.94 | 100% | 59% | 0.87 | 100% | 41% | 0.31 | 24% | 5% | 0.39 |
| **Setting-II** | 49% | 11% | 0.96 | 100% | 61% | 0.93 | 100% | 39% | 0.40 | 25% | 5% | 0.66 |
| **Setting-III** | 100% | 39% | 0.98 | 100% | 69% | 0.94 | 100% | 40% | 0.40 | 45% | 8% | 0.64 |

*NPV = negative predictive value; PPV = positive predictive value, % of probs = the percentage of all probabilities, % of pts = the percentage of patients that had probabilities for at least one disease below or above the given cut-off.*

Finally, our main interest was whether G-Prob has the ability to correctly classify disease status. To test this, we depicted receiver operating characteristic (ROC) curves and summarized the total performance with the area under the curve (AUC) for G-Prob with disease match. The overall discriminatory capacity of G-Prob was highly accurate with an AUC 0.86 [95%CI 0.86-0.86] (Fig. 2C, an overview of AUCs for the individual diseases is provided in **Table S6**). The depiction of the precision (PPV) versus the recall (sensitivity) is provided in the supplement **Fig. S12**).

The simulation showed that genetic information was substantially different for the five diseases and could have the potential to prioritize diagnoses. However, the genetic architecture employed in simulations may differ in real patient data, due to differences in reported effect sizes and unappreciated environmental factors. Therefore, we tested how well genetic information separated patients in real data in three settings. For this we collected patients from biobank cohorts in which genetic data were linked to electronic medical records.

Setting I comprised of patients from eMERGE (Electronic Medical Records and Genomics), a network that has amassed clinical data and genome-wide genotyping data from 12 healthcare networks from throughout the United States (38). This consortium includes medical centers with biobank genetic information linked with electronic medical records (EMR) data, such as diagnosis (International Classification of Diseases and Related Health Problems [ICD]-9 and −10)billing codes. Among 83,717 subjects in eMERGE, 72,624 individuals were self-described as white, and of those 52,623 were not included in setting II and III (Partners Biobank). After testing the performance of the number of ICD codes for case identification with EMR-reviewed data as the gold standard (**Fig. S2**), we chose a cut-off of ≥20 ICD9/10 codes for each disease to classify patients by their eventual true diagnoses. Hereby, we identified 1,211 patients with one of 5 diseases (**Fig. S3** and **Table S3**).

Reflecting the uncertainty of billing code-based diagnoses, G-Prob probabilities were well calibrated with disease status but slightly lower (beta=0.90) (Fig. 2B). We observed that 13% of the G-Prob probabilities were ≤0.05, corresponding to an NPV=0.94. Using this cut-off, we could identify and classify ≥1 of the 5 diseases as highly unlikely in 619 (51%) patients, ≥2 diseases in 135 (11%) patients and ≥3 diseases in 10 (<1%) patients. For 24% of patients, there was a single disease with ≥0.5 probability, corresponding to 5% of the total assigned probabilities. In this data set, this corresponded to a PPV of 0.39. G-Prob showed a modest AUC discriminating between diseases (AUC=0.69 [95%CI 0.67-0.71], Fig. 2C).

Since misclassification due to using ICD-codes may have influenced setting I, in setting II we collected patients through the most precise method of patient selection: manual review of patients’ complete medical record. This was possible in the data from the Partners HealthCare Biobank, which comprises of >80,000 subjects from Boston based hospital centers recruited from approximately 1.5 million patients(30). The samples are linked to clinical EMR data. From the 12,604 self-described white Biobank patients with genotype data available, we pre-selected potential patients using a simple algorithm combining ICD-codes and clinical characteristics (**Fig. S4** and **Table S4**). Then, a rheumatologist reviewed the medical records manually for clinical criteria as formulated by the American College of Rheumatology and/or the European League Against Rheumatism (14–21). We identified 245 patients with one of the five diseases of interest.

The more stringent patient selection in setting II resulted in a greater correspondence between G-Prob and disease match (beta=1.08) (Fig. 2B). Here, 11% of the probabilities were ≤0.05, corresponding to an NPV=0.96. At this cut-off, it is possible to de-prioritize ≥1 disease in 119 (49%) patients, ≥2 diseases in 13 (5%) patients and ≥3 diseases in 3 (1%) patients. Furthermore, 25% of the patients had a single disease with a G-Prob ≥0.5 (Table 1), which was 5% of all calculated probabilities. A G-Prob ≥0.5 corresponded to a PPV=0.66. We observed that the accuracy was closer to the accuracy in the simulation data with AUC=0.81 [95%CI 0.76-0.84](Fig. 2C).

To ensure that one strong genetically-determined disease did not skew the results, we conducted several sensitivity analyses (**Supplement 2).** When we calculated the AUC for each individual disease, G-Prob showed similar performance for the individual diseases (**Fig. S7**). Rerunning G-Prob five times excluding one disease each time resulted in similar AUC values (**Fig. S8**).

Since our shrinkage factor of 0.5 was arbitrarily chosen, we tested G-Prob’s calibration with disease outcome, the log likelihood and entropy score (39) using different shrinkage factors for the odds ratios (ORs) in the GRS and found that the results did not substantially differ (**Fig. S9**).

We grouped peripheral SpA(17) with axial SpA(18, 19) but separated PsA(20) as a different disease phenotype, since there is extensive GWAS data on axial SpA and PsA (17–20). We tested whether the inclusion of peripheral SpA cases had influenced the result by rerunning analyses without those cases. The results remained similar (beta 1.09 (95%CI 1.01-1.18), AUC 0.81 (95%CI 0.78-0.84)).

As G-Prob was developed to discriminate patients presenting with similar symptoms, we tested this exact hypothesis in setting III by manually reviewing the records of all patients that received ICD codes from a rheumatology clinic for the presence of inflammatory arthritis at their first visit. If a patient had inflammatory arthritis, we (again manually) reviewed the complete record until the final diagnosis. From the 1,808 Partners Biobank patients that visited a rheumatology outpatient clinic, 282 had inflammatory arthritis without a previous diagnosis. We excluded seven cases in which the patient did not fulfill any classification criteria, but the rheumatologist did diagnose the patient with one of the five diseases, because we could not determine whether the medical record lacked information, or the patient was misclassified by the rheumatologist. Of the remaining patients, 79.4% were diagnosed using complete follow-up information with one of the diseases of interest (**Fig S5** and **Table S5**), which is similar to past studies(40, 41). We classified the remaining 20.6% as “other diseases” and included this as a sixth disease group in G-Prob, accounting for the fact that patients with inflammatory arthritis can have conditions in addition to the five most common diseases. In addition, we used the diseases’ prevalence within the outpatient clinic. Due to these two adjustments, made possible by the prospective design, G-Prob reflected the most realistic disease risk. We excluded RA patients without anti-CCP antibody-status documented, resulting in 243 patients for analysis.

Again, G-Prob probabilities calibrated well with real disease status (beta=0.99). Here, 39% of the G-Probs were ≤0.05. At this threshold, we could de-prioritize ≥1 disease in 100% of patients, ≥2 diseases 203 patients (84%), ≥3 diseases in 98 patients (40%) and ≥4 diseases in 27 patients (11%), based on genetic data alone. Furthermore, 45% of the patients had a single disease with G-prob≥0.5; representing 8% of the calculated probabilities (PPV=0.64). The discriminatory ability was similarly high as in previous settings (AUC=0.84 [95%CI 0.81-0.86]) (Fig. 2B&C).

Since the rheumatology clinics at Partners Healthcare are tertiary referral centers with an interest in RA, this setting was enriched for RA patients. Hence, the high difference in disease prevalences influenced the results in setting III. Sub-analyses assigning uniform prevalences showed an improved performance for the less prevalent diseases (**Fig. S10**).

The prospective patient identification enabled us to compare G-Prob’s performance with the diagnosis of the rheumatologist at a patient’s first visit. Compared with the final diagnosis after complete follow-up (median 7 years), we observed that 35% of the patients were misclassified by their rheumatologist at their first visit (Fig. 3A). The misclassification compared to eventual diagnosis was 35% for RA, 33% for SLE, 50% for SpA, 42% for PsA, 11% for gout and 37% for other diseases. Of these initial misdiagnoses, 43% had a genetic probability ≤0.5, 29% ≤0.2 and 9% ≤0.05. The difference of G-Prob’s probabilities between the eventual diagnosis and the incorrect initial diagnosis, favored the probability of the eventual diagnosis (Fig. 3B): for 65% of the patients G-Prob’s probability of the correct disease was higher than G-Prob’s probability of the initial diagnosis of the rheumatologist.

**Fig. 3.**
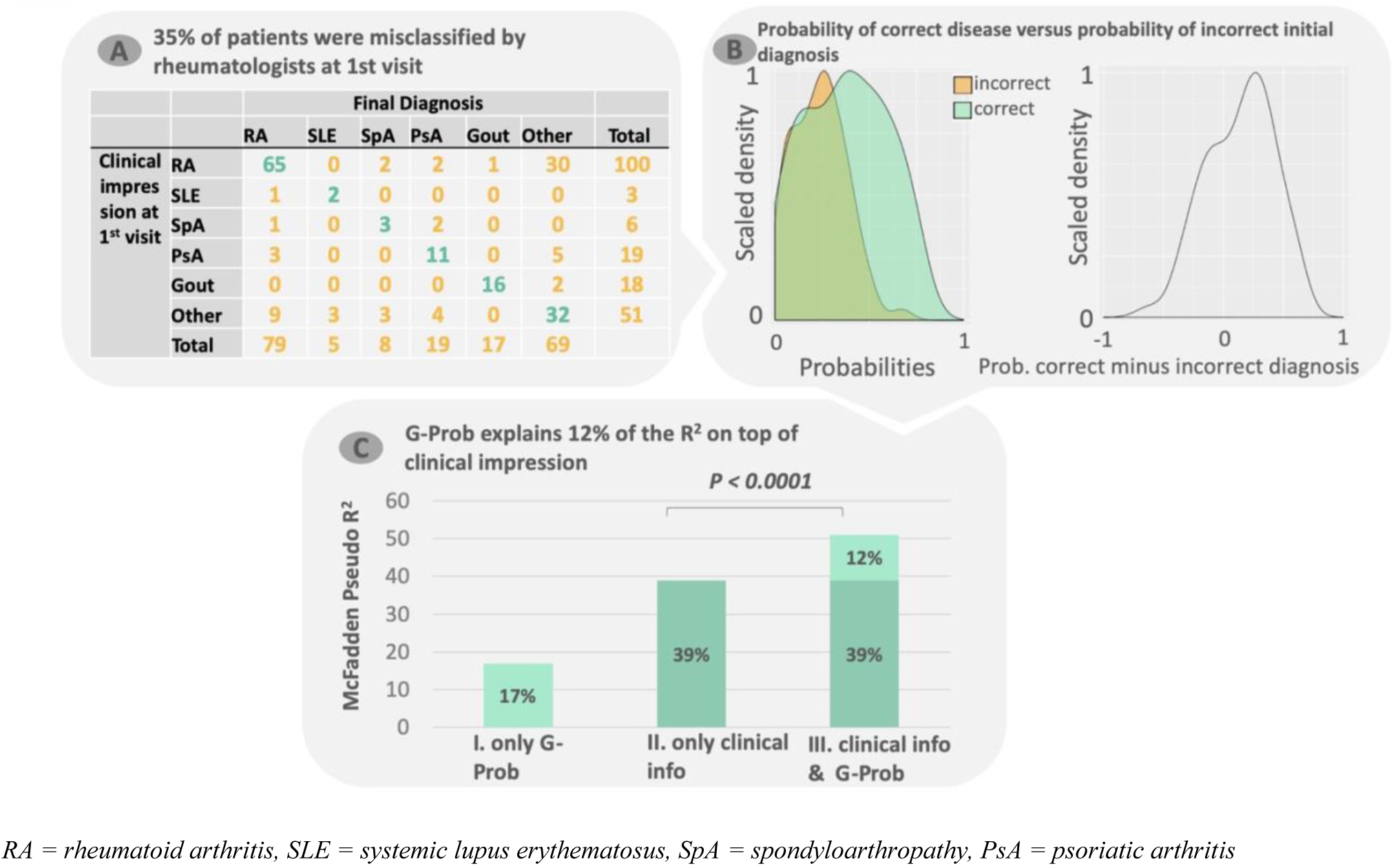
Value of G-Prob in addition to clinical information at first visit. In setting III, we collected the (highest ranked) diagnosis of the rheumatologists at the first visit as proxy for all the clinical information and matched this with the final diagnosis (box A). 35% of these initial diagnoses were different from the final diagnosis and are thus misclassifications. To test if G-Prob could have guided the rheumatologist towards the correct diagnosis, box B depicts the density of patients’ probabilities of the real (correct) disease (green) and the probabilities of the incorrect initial (clinical) diagnosis from table A (orange) (left panel) as well as the difference between the two (right panel). Here, 65% of the probabilities of the correct disease were higher than of the incorrect clinical diagnosis In box C. the graph shows the McFadden’s R_2_ on the y-axis of three multinomial logistic regression models: I) Model with the G-Prob only II) Model with only clinical information (rheumatologist diagnosis at first visit) III) Model with both clinical information and G-Prob.

Taking the highest G-Prob from each patient, 53% of those corresponded with the final diagnosis. This was 77% for the top 2 highest probabilities and 87% for the top 3. The McFadden’s R_2_ (which is the logistic regression equivalent of the explained variance of linear regression analyses)(42) of G-Prob alone was 17% and 39% for the diagnosis at first visit. Adding G-Prob to the clinical information significantly improved the model with an increase of 12 percent points yielding a R_2_ of 51% (*P*<0.00001) (Fig. 3C). Thus, the availability of G-Prob’s information would have improved the rheumatologist’s differential diagnosis.

Since rheumatologists can improve their differential diagnosis with readily available serologic testing, we explored whether G-Prob serologic data could improve clinical insight when added to our model. Serologic data added to clinical data improved R_2_ by 22%. Serologic data added to clinical + genetic model improved R_2_ by 12%. Since several studies describe a strong association between genetics and serology(43–45), we were surprised to find that serologic testing improved the model’s accuracy in addition to the genetic information. To model a setting where serology is available at the first visit, even with clinical + serologic data present, G-Prob significantly improved the R_2_ by 12 percent point (**Table S7**).

## DISCUSSION

The number of patients with DNA genotyped prior to their first visit to a clinic is rapidly expanding. We investigated whether genetic information can help clinicians initially prioritize likely diagnoses and deprioritize unlikely ones among patients presenting with inflammatory arthritis. We observed that genetic information adds value to the clinical information obtained at the initial encounter, even when serologic data are available. Taken together, pre-existing genetic data may can be considered part of a patient’s medical history with the potential to improve precision medicine in the modern outpatient clinic.

Our results demonstrate that genetic data can provide probabilistic information to discriminate between multiple diseases presenting with similar clinical signs and symptoms. We investigated inflammatory arthritis, a hallmark of many rheumatic diseases associated 80% of the time with the diseases that we focused on in our study: RA, SLE, SpA, PsA and gout(14–21). Due to the evolution of symptoms over time, disease classification by a clinician at a first visit is challenging, evidenced by the 35% misclassification of patients by the rheumatologist at first visit. Our data strongly suggest that using increasingly available genetic information would reduce this misclassification rate.

Genetic risk scores (GRSs) have been studied for both prediction of rheumatic disease progression(5–7) and for disease susceptibility(8–11). These GRSs had modest predictive value in determining case versus control status. Given the low prevalence of rheumatic diseases, most tests perform poorly on a population level since the pre-test disease probability is low(46). As inflammatory arthritis is not present in healthy individuals, symptom-based selection substantially increases the pre-test probability for disease, resulting in an increased post-test probability that renders probabilistic predictions useful in the clinical setting.

While G-Prob’s magnitude was consistent with real disease risk and had a high AUC in simulated data, it also performed well in data in which clinical diagnoses were obtained from EMRs. Notably, performance was better in rigorously reviewed cases.

Our study contained two settings with complete manual review of patients medical records. In setting II, by preselecting records with simple disease-specific algorithms validated by medical record review, each disease was well represented, and we were able to test the performance of G-Prob for the individual diseases. G-Prob’s performance did not significantly differ between the individual diseases compared to the overall performance.

In settings I and II, we selected patients based on criteria aiming for well-defined cases (see **Fig. S3** and **S4**). Here patients with less severe disease may have been excluded. It is likely that patients with less severe disease have less genetic risk and that including such patients would decrease the value of G-Prob. Therefore, we consider setting III particularly interesting. In this setting, we selected patients who presented with inflammatory arthritis for the first time at a rheumatology clinic, by reviewing the records of all patients that received ICD codes from a rheumatology clinic. If a patient had inflammatory arthritis at their first visit, we followed their disease course until the final diagnosis. This thorough patient selection ensured that the analysis most optimally represented the study hypothesis that genetic data can facilitate disease differentiation in patients with early symptoms. Although this setting had a relatively low patient number and had an extra level of complexity due to the addition of the category “other diseases”, G-Prob was able to differentiate the disease categories in patients presenting with inflammatory arthritis showing similarly good results as in setting II.

We note that G-Prob’s performance depends on the availability of genetic data for different diseases. In our case, genetic data from genome-wide association studies of rheumatic diseases were primarily available from white/European individuals. Lack of diversity of genetic studies is potentially crippling the clinical applicability of G-Prob and other genetic risk score strategies(47). We note there is a rapid explosion of biobank data, where patients with different backgrounds and a wide range of phenotypic information are being obtained alongside genetic data. These studies, such as the UK Biobank (48), may offer more clear data on the differences and similarities in the genetics of different inflammatory arthritic conditions, such as axial and peripheral SpA. Sequencing of large cohorts may reveal effects of unidentified variants that could improve the performance of G-Prob. In our study, we intended to test the performance of genotypes obtained with standardized technology that can be readily available when EMRs and genetic data are integrated. The advantage of this approach is that it makes G-Prob easily implementable and transformable to other diseases as one only needs the risk variants as described in previous genome-wide association studies.

Although clinicians have several tests to prioritize their differential diagnosis, complex diseases such as rheumatic disease take notoriously long to diagnose; a third of the arthritis patients are initially classified as undifferentiated even when all tests results are available (49), and 48% of SLE patients have to wait >6 months to receive their diagnosis (50). We expect that in the future an increasing number of patients will have genetic data available prior to their visits, making it possible for a clinician to request a G-Prob calculation while preparing patients’ visit (while reviewing a patients’ medical history before their visit). When more information is known about a patient disease (for instance at later visits), the value of G-prob will inevitably decrease. We have not tested how much it decreases over time. Since we found that G-Prob’s significantly contributes to disease classification even when serologic data is present (**Table S7**), we conclude that G-Prob can complement the arsenal of tests available to future clinicians.

Non-genetic information available at a first visit such as family history and age could further improve the performance of predictive tests such as G-Prob. Future studies could combine both genetic and non-genetic factors aiming for a more precise prediction model. We restricted our study to genetic factors since we designed a proof-of-principle study of the value of genetics.

The importance of differentiating diseases in patients with similar presenting symptoms is relevant not only in rheumatology, but in many other clinical settings in which patients present with similar symptoms: endocrinology (late onset type 1 diabetes versus type 2 diabetes), pulmonology (COPD vs asthma), cardiology (subsets of heart failure) etc. Numerous genetic risk factors are known for such diseases, but not yet translated to clinical practice. The G-Prob method can serve as a template to develop symptom tailored disease-differentiating tests.

Though we have tested G-Prob extensively, the current study does not give a specific threshold by which G-Prob would accelerate diagnostic care. With our current proof-of-principle study, we have demonstrated that genetic information is able to improve the differential diagnosis, but whether this would translate into an acceleration of the diagnostic process requires a targeted comparative prospective study where clinicians have G-Prob data for a selection of their patients.

## MATERIAL AND METHODS

### Study populations

#### Validation through simulation

To validate G-Prob in an optimal setting, we simulated a population of 1 million samples using the population-based allele frequencies of uncorrelated risk SNPs as reported for European samples(51). First, we calculated the number of samples with a hetero- or homozygotes (major or minor) genotype using the reported minor allele frequency (MAF) of Europeans from Ensembl(52), ensuring an allele distribution as expected under the hardy-weinberg-equilibrium:

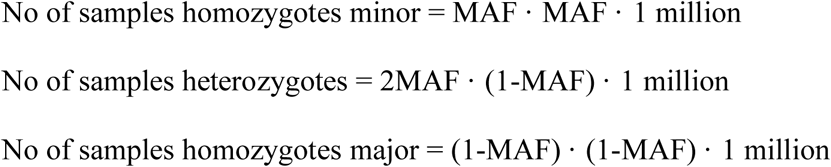

Then, we randomly assigned a SNP genotype (0, 1, or 2 alleles) to each sample by sampling without replacement and repeated this for all non-HLA SNPs. For the HLA region, we randomly sampled complete profiles from Jia et al.(53) reference panel to ensure that the strong linkage structure of the HLA region would be present in our simulated data.

Next, for each sample the population-based disease probability was calculated (see “G-Prob development” *Equation 3*) for each of the five rheumatic diseases. Case status was assigned when a patient’s disease probability was higher than a randomly sampled number between 0 and 1 from a binominal distribution. Using this approach, we obtained 49,151 patients having one of the five diseases (**Fig. S1**). In this set of 49,151 patients, we calculated the within patient risk for each disease (G-Prob) (*Equation 4 and 5*).

#### Setting I: ICD based patient selection

eMERGE (Electronic Medical Records and Genomics) Network(38) is a consortium of medical centers with electronic medical records (EMR) linked with genetic information and Biobank data such as the billing diagnosis codes (codes for International Statistical Classification of Diseases and Related Health Problems (ICD)). This includes 83,717 patients from 12 medical centers. The eMERGE network is an NIH approved collaboration. We obtained consent from the network to use the imputed genotypes, self-reported ethnicity and ICD9 and ICD10 data for this study. We selected patients with one of our five rheumatic diseases using relevant ICD codes.

Previous studies have found that disease specific ICD-codes are registered for patients even if they do not have that disease (54, 55). To find the optimal number of ICD codes for patient selection, we explored the ICD code performance in previously reviewed medical records for rheumatoid arthritis in the Partners Biobank(56). Here, one rheumatologist screened the medical records, and assigned case status both by the ACR2010 criteria(14) as well as her expert opinion. We used this data as the golden standard and checked the test characteristics of different thresholds of ICD9 codes aimed to identify the RA patients (**Fig. S2 and Table S2**). Also in our data, we found that one ICD code does not serve well to identify cases. Based on this analysis, we decided to select patients in the eMERGE dataset if they had >20 ICD9 or ICD10 codes for one particular disease of our five diseases of interest (**Fig. S3**). This cutoff ensured us that the patient status was highly likely, though it might have excluded patients that had less ICD9/10 codes because they were recently diagnosed or because they did not frequently visit the clinic. **Table S3** summarizes the patient characteristics of this dataset.

#### Setting II: Patient selection through ICD-codes and manual review of medical records

Partners HealthCare Biobank comprises of >80,000 subjects from Boston based hospital centers (Brigham and Women’s Hospital, Massachusetts General Hospital, Faulkner Hospital, Newton-Wellesley Hospital, McLean Hospital, North Shore Medical Center, and Spaulding Rehabilitation Network) recruited from approximately 1.5 million patients(56). Written consent was obtained from each patient before their data was included in the Biobank. We obtained approval from the Partners Institutional Review board (IRB) to use the genotypes of these patients and access their clinical records.

Before review of the medical records, we preselected records on clinical features that are typical for rheumatologic diseases (**Fig. S4**). The selected records were reviewed manually (**Fig. S6**) and patients were identified by applying the disease classification criteria(14–21). **Table S4** summarizes the patient characteristics of this dataset.

#### Setting III: Prospective patient collection

In setting III, we collected patients in a prospective manner by manually reviewing the records of all patients that received ICD codes from a rheumatology clinic on the presence of inflammatory arthritis at their first visit. We performed a complete (again manual) review on the patients with inflammatory arthritis until their most recent visit. This setting consisted of patients collected from the same biobank as setting II. In contrast to the preselection of medical records based on case specific features in setting II, in setting III, we examined all medical records starting at the first visit to the rheumatology clinic of Biobank patients (n=1,808) to identify patients who presented with inflammatory arthritis to a rheumatologist for the first time. Subsequently, we reviewed the medical records for the criteria-based diagnoses until the present time (**Fig. S6**). Patients that did not fulfill any of the criteria of our five diseases of interest were classified as “other diseases”. We also registered the (highest ranked) diagnosis made by the rheumatologist at the first visit. See **Fig. S5** and **Table S5** for detailed patient (selection) description.

### G-Prob calculation

#### General principle of G-Prob

Genome-wide-association studies (GWAS) have used logistic regression to identify disease susceptibility factors in patients by comparing them with healthy controls. The resulting odds ratios (OR) refer to a relative increase in population-based odds. Our genetic probability model, G-Prob, uses these log odds ratios (logOR) to create a weighted genetic risk score (GRS)(57). G-Prob consists of two steps (Fig. 1B):

#### G-Prob step 1

The genetic risk score of individual *i* for disease *k* is defined as:

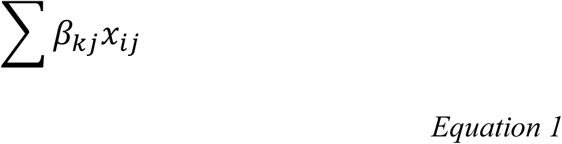

With

- *x_ij_* the number of risk alleles of SNP j present for individual *i*.
- *β*_*kj*_ the logOR for SNP *j* obtained from previous GWAS studies for disease *k*
- To correct for possible overestimation of the effect sizes due to publication bias (58), we shrank the logORs of each genetic variant by multiplying them with 0.5.

The genetic risk score can be used to calculate for each subject a population level disease probability (*P_ki_*) of each patient *i* for each disease *k*, using the logistic regression formula:

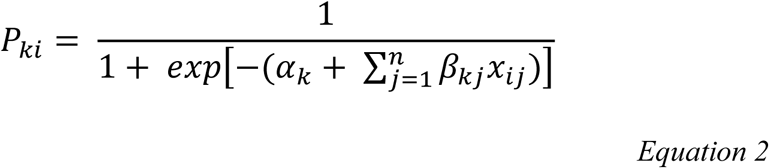

The GRS of each disease is combined with an intercept that ensures the mean probability is equal to the predefined disease prevalence. Where *α*_*k*_ is an unknown constant, which is estimated by assuming that the mean predicted pre-test probability is equal to the population prevalence of disease *k*, i.e. by minimizing the following estimation equation:

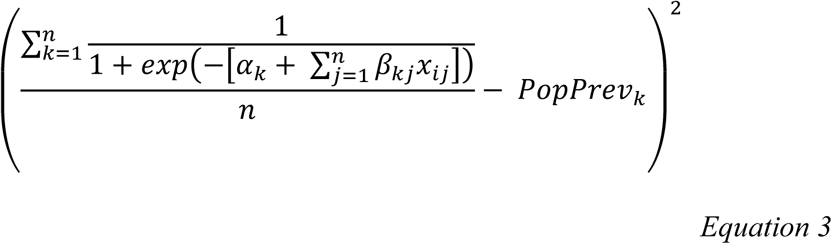

Where *PopPrev_k_* is the assumed prevalence for disease *k* in the general population.

#### G-Prob step 2

The next step is to calculate the conditional probabilities of the diseases of interest *k*, given that the patient has one of the 5 diseases (Pr(Y*_k_*=1| Σ(Y*_k_* = 1)). These within patients’ genetic probabilities (*cP_ki_*) of each patient (*i*) for each disease (*k*) were obtained through normalization of the population risk:

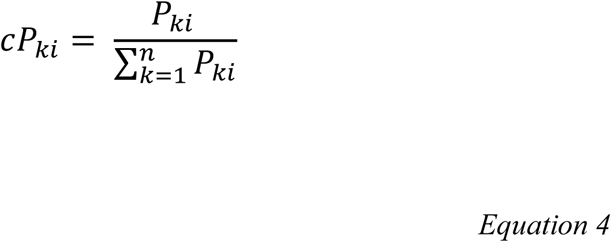

The G-Prob gives each patient a probability for each of the five diseases of our interest. The output of G-Prob is visualized in Fig. 1A. The final product of G-Prob per patient is as follows:

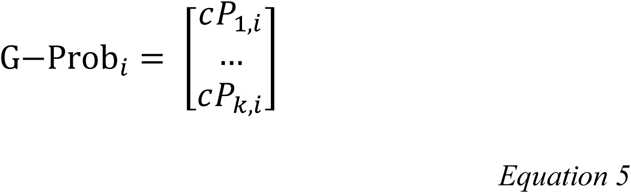

To summarize, G-Prob assigns a disease probability for each disease of interest based on a weighted genetic risk scores for the diseases taking into account a predefined (sex specific) disease prevalence assuming a patient has one of the possible diseases. G-Prob can be used to discriminate between any set of diseases providing there is sufficient knowledge of the genetic risk.

#### Derivation of G-Prob in the rheumatology clinic

In our G-Prob, we included the five most common inflammatory arthritis-causing diseases in G-Prob: rheumatoid arthritis (RA), systemic lupus erythematosus (SLE), spondyloarthropathy (SpA), psoriatic arthritis (PsA) and gout.

#### Selecting the prevalences

For the non-prospective collected samples (simulation and setting I and II), we assumed a population prevalence of 1% for each disease. To incorporate the genetic variant sex, we used sex-specific prevalences using the sex specific risk ratio. For example, the risk for RA is 3 times higher in women than in men, and assuming an overall prevalence of RA of 1%, and a 1:1 ratio of women and men, the RA prevalence is 1.5% in women and 0.5% in men. So, equation 3 was minimized for men and women separately. We used F:M risk ratio of 3:1, 11:1, 0.3:1,1:1, 0.3:1, for RA, SLE, SpA, PsA, and gout respectively.

#### Derivation of the GRS in the rheumatology example

We took genetic risk variants that obtained genome-wide significance (p<5×10_-8_) for one or more of the five diseases from Immunobase(51) or (when not available) from the most recent genome wide study on people from European decent. We used the diseases specific OR. So, SNPs that contribute to the susceptibility of several rheumatic conditions had different ORs for each disease. If a SNP was not associated with the other diseases, the ORs for those diseases was 1.0. If there were two SNPs associated to the same disease in linkage with each other (r_2_> 0.8) we selected the SNP with the highest OR. For gout the reported risk effects were beta’s per uric acid increase. Köttgen et al have developed a prediction model for gout translating the genetic beta’s for uric acid levels into risk groups and assigned an OR to each group(23). We used this risk group categorization and the corresponding OR to calculate the genetic risk score (GRS) in gout.

For HLA, we searched recent large studies that tested the HLA variants dependently. This ensured that the ORs used in our study were corrected for the strong linkage disequilibrium in this region. Since the HLA risk differs between CCP+ and CCP-RA, we created a separate GRS for CCP+ and CCP-RA using their specific HLA ORs in the datasets where CCP-status was available (simulation and setting II and III). Before step 2 of G-Prob, we combined CCP+ and CCP-patients into one population probability for RA for each patient. In the settings where no CCP status was known, we used the HLA variants for CCP+ RA.

The final G-Prob model consisted of 208 SNPs outside the HLA region and 42 HLA variants such as SNPs, haplotypes and alleles. The number of variants for each disease ranged from zero HLA variants (gout) to 21 variants (CCP+ RA) and 18 non-HLA variants (PsA) till 93 variants (CCP+ RA). **Table S1** (TableS1_G_Prob_risk_variants.xlsx) summarizes the included risk variants’ information.

#### Optimizing G-Prob in the prospective setting III

In the prospective analysis (setting III) the category “other diseases” was added as a sixth category to the five diseases. The ORs for the GRS of the “Other” group were all 1.0, as thus far no data on the genetic risk profile of these patients is available. Through the prospective design, we also obtained the real outpatient-clinic prevalences which we incorporated into the G-Prob calculation.

### Data analysis

All analyses were performed in R version ≥3.2(59).

Our primary test was the overall performance of G-Prob combining all diseases. G-Prob gives each patient a probability for each disease of interest. Only one of these diseases is a patient’s real disease. For analyses on G-Prob’s overall performance, we grouped all probabilities of all patients into one vector and created a corresponding vector indicating whether a particular probability corresponded with a patient’s real disease which we call disease match. Hereby the binary disease match vector contained 20% instances where the probability of one disease matches a patient’s real disease match and 80% non-match. If G-Prob is able to differentiate diseases, the real disease matching probabilities would be higher than the non-matching probabilities.

We tested the performance of G-Prob as follows.

a. First, we examined how well G-Prob was calibrated with disease outcome. We performed a linear regression model without intercept using probabilities as independent variable and disease match as dependent variable. The regression coefficient (beta) describes how well the model was calibrated (ideally beta is one)(60).
b. Second, we explored the ability of G-Prob to correctly classify disease status. For this we created receiver operating characteristic (ROC) curves and summarizes the total performance with the area-under the curve (AUC) of multi-class classifications(61). The ROC depicts the true positive rate (sensitivity (y-axis)) against the false positive rate (1 minus specificity, x-axis) for the probabilities as test-variable and disease match as gold standard. Higher AUCs indicate better classification. A random predictor would have an AUC of 0.5, while a perfect predictor would have an AUC of 1. Given our two vector data summary, our AUCs are so-called microAUC as described in the R Package multiROC(62). In the supplement, we also provide the macroAUC which averages of the AUCs of each disease group. The AUC is 1 for a perfect predictor. We obtained 95% confidence intervals by bootstrapping (100 resamples). We compared the AUCs (supplement data) using DeLong’s method within R package, pROC(61).

We corrected the ORs in the GRS with an arbitrarily choosing shrinkage factor of 0.5. To test if this choice has influenced our results, we recalculated the G-Prob with 10 different shrinkage factors between zero and one. For each group of G-Probs we tested the calibration, the log likelihood (a measure of the how well a model fits the data where the higher the log likelihood, the better the model fits) and the entropy scores (a measure for the disorganization of values, where the lower the entropy the more organized the values are) :

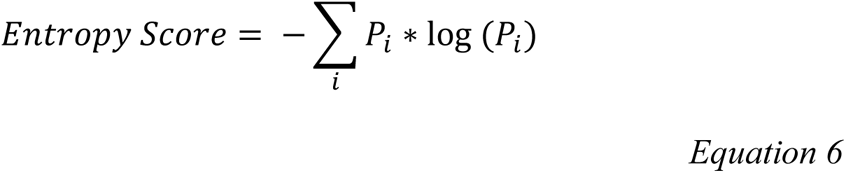

To test G-Prob’s clinical performance in setting III, we used the rheumatologists’ diagnosis at first visit as a proxy for the clinical information. We applied multinomial logistic regression with the six disease categories as the dependent variables and the diseases’ probabilities as well as the clinical information as independent variables. We calculated the McFadden’s pseudo-R_2_ (the logistic regression equivalent of the explained variance of linear regression analyses)(42) (*Equation 7*) to compare how different models (I) the G-Prob; (II) the clinical information; (III) the combination of the clinical information with the G-Prob, predict. With the likelihood ratio test, we assessed if the addition of the G-Prob to the clinical information significantly improved the model.

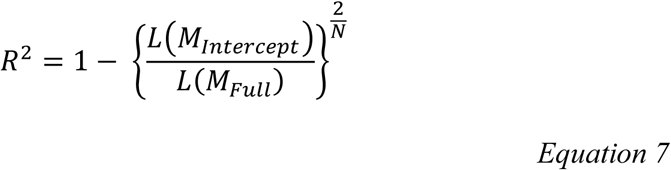

#### Genotyping and imputation

##### 1. Setting I

The eMERGE samples were genotyped on 78 Illumina and Affymetrix array batches with different genome coverage(29). Missing genotypes were obtained by imputation using the Michigan Imputation Server with the minimac3 imputation algorithm(54) and the HRCv1.1 reference panel(63). Before imputation, data was curated using PLINK version 1.9b3 (64) applying the following settings: maximum per-SNP missing of <0.02, maximum per-person missing of <0.02, minor allele frequency of >0.01 and Hardy-Weinberg disequilibrium p-value (exact) 0.00001.

We required each genetic variant to have an imputation quality ≧0.8. In case a variant of interest was not present in the post-imputed post-curated data, we searched for a proxy that was in close linkage with the original variant (r_2_>0.8).

##### 2. Setting II and III

At time of the initiation of this study, 15,047 patients of the Partners Biobank were genotyped in three consecutive rounds. The first set of 4,930 patients were genotyped on the Illumina Multi-Ethnic Genotyping Array (MEGA) chip. The second and third set were genotyped on Illumina Multi-Ethnic Genotyping Array Expanded (MEGA Ex) chip(65).

To obtain all relevant SNPs for the GRS, we imputed the missing genotypes on the Michigan imputation server with the minimac 3 imputation algorithm and 1000genome phase 3 v5 as reference panel. Before imputation, data was curated using PLINK version 1.9b3(64) applying the following settings: maximum per-SNP missing of <0.02, maximum per-person missing of <0.02, minor allele frequency of >0.01 and Hardy-Weinberg disequilibrium p-value (exact) 0.00001.

We required each genetic variant to have an imputation quality ≧0.8. In case a variant of interest was not present in the post-imputed post-curated data, we searched a proxy that was in close linkage with the original variant (r_2_>0.8).

##### 3. HLA imputation

Both for eMERGE and the Partners HealthCare data, we imputed the HLA region using SNP2HLA(57). This tool imputes amino acid polymorphisms and single nucleotide polymorphisms in HLA within the major histocompatibility complex (MHC) region in chromosome 6. The T1DCG v1.0.3 reference panel was used(53).

## ACKNOWLEGMENTS

Conflict of interest: none

Funding: NIH/NHGRI U01HG008685, ReumaNederland 15-3-301, Harold and DuVal Bowen Fund, NIH p30 AR072577

