## Supplemental Figures and Tables for "Using genetics to differentiate patients with similar symptoms: application to inflammatory arthritis in the rheumatology outpatient clinic"

### Supplementary Online Content

**Supplement 1** Patient selection

#### 1. Validation: Simulation

**Fig. S1** Flow simulation study

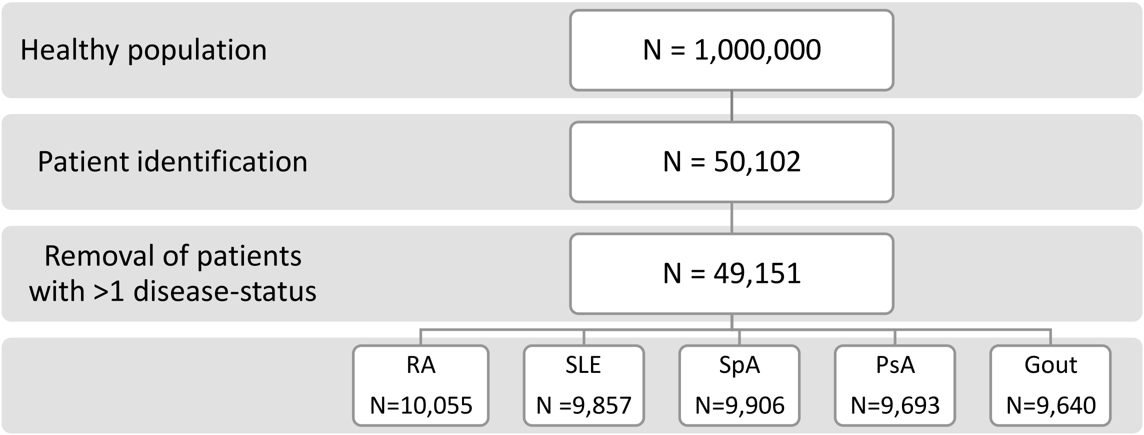

#### 2. Setting I: ICD based patient selection

**Fig. S2** Test characteristics of different ICD9 cut-offs for identification of rheumatoid arthritis cases using the reviewed medical records data as the golden standard.

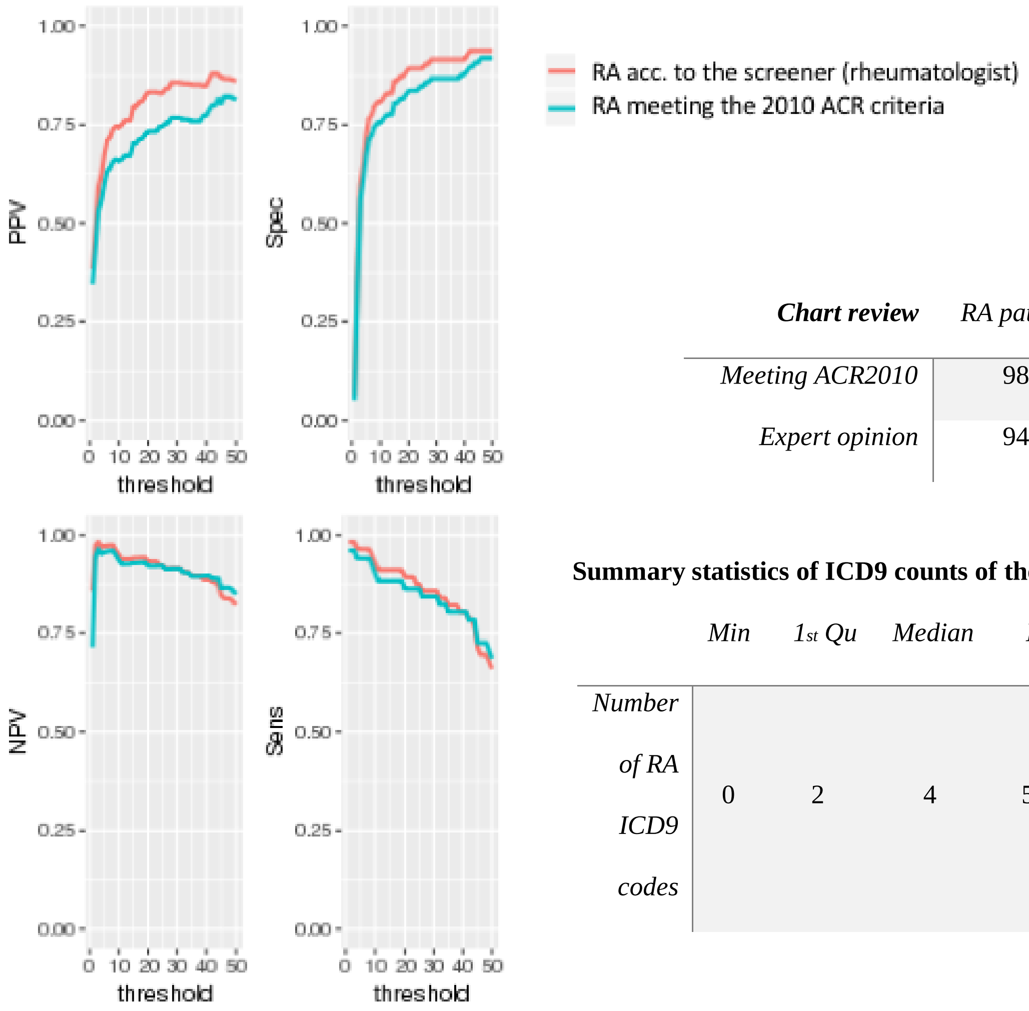

| ***Chart review*** | *RA patients* | *RA non patients* |
|---|---|---|
| *Meeting ACR2010* | 98 | 51 |
| *Expert opinion* | 94 | 56 |

**Summary statistics of ICD9 counts of the reviewed notes**

| | *Min* | *1st Qu* | *Median* | *Mean* | *3rd Qu* | *Max* |
|---|---|---|---|---|---|---|
| *Number of RA ICD9 codes* | 0 | 2 | 4 | 54.9 | 54 | 1315 |

**Table S2** ICD9 and ICD10 codes used to identify patients in the eMERGE dataset setting I

| Phenotype | Code type | |
|---|---|---|
| *RA* | ICD9 | 714.0, 714.1, 714.2, 714.81 |
| | ICD10 | M05.00, M05.10, M05.141, M05.19, M05.20, M05.212, M05.271, M05.29, M05.30, M05.39, M05.442, M05.50, M05.59, M05.60, M05.621, M05.641, M05.642, M05.661, M05.69, M05.70, M05.711, M05.712, M05.719, M05.721, M05.722, M05.729, M05.731, M05.732, M05.739, M05.741, M05.742, M05.749, M05.751, M05.752, M05.759, M05.761, M05.762, M05.769, M05.771, M05.772, M05.779, M05.79, M05.80, M05.821, M05.822, M05.831, M05.832, M05.841, M05.842, M05.849, M05.861, M05.862, M05.871, M05.872, M05.89, M05.9 , M06.00, M06.011, M06.012, M06.021, M06.022, M06.029, M06.031, M06.032, M06.039, M06.041, M06.042, M06.049, M06.051, M06.052, M06.059, M06.061, M06.062, M06.069, M06.071, M06.072, M06.079, M06.08, M06.09, M06.1, M06.262, M06.271, M06.30, M06.321, M06.322, M06.332, M06.341, M06.342, M06.349, M06.371, M06.39, M06.4, M06.80, M06.812, M06.821, M06.822, M06.831, M06.832, M06.841, M06.842, M06.849, M06.851, M06.852, M06.861, M06.862, M06.871, M06.872, M06.879, M06.88, M06.89, M06.9 |
| *SLE* | ICD9 | 710.0 |
| | ICD10 | M32.0, M32.10, M32.11, M32.12, M32.13, M32.14, M32.15, M32.19, M32.8, M32.9 |
| *SpA* | ICD9 | 720.0, 720.1, 720.2, 720.8, 720.81, 720.89, 720.9 |
| | ICD10 | M45.9, M46.00, M46.1, M49.80, M46.80, M46.90 |
| *PsA* | ICD9 | 696.0 |
| | ICD10 | L40.52, L40.51, L40.50, L40.59, L40.54 |

| | | |
|---|---|---|
| *Gout* | ICD9 | 274, 274.0, 274.00, 274.01, 274.02, 274.03, 274.1, 274.10, 274.11, 274.19, 274.8, 274.81, 274.82, 274.89, 274.9 |
| | ICD10 | M10.00, M1A.9XX0, M1A.00XX1, M10.30, M10.9, M10.40 |

**Fig. S3** Flow patient selection in setting I

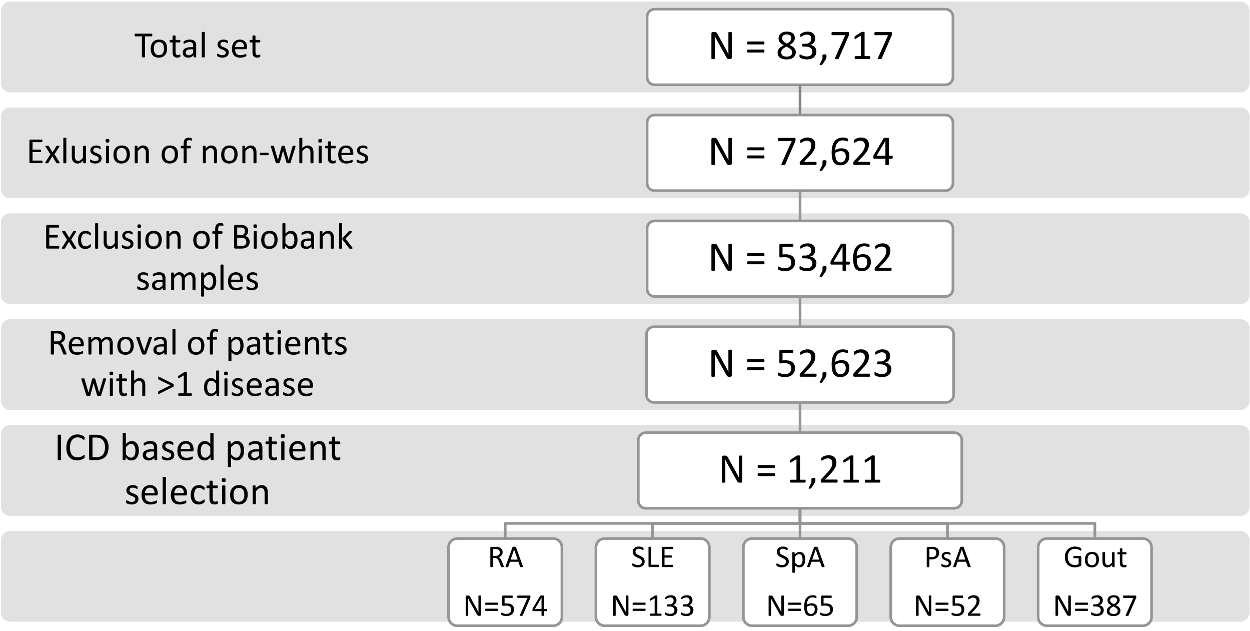

**Table S3** Patient characteristics setting I

| | **Patients included in this study** | | | | | |
|---|---|---|---|---|---|---|
| | RA | SLE | SpA | PsA | Gout | Total patients |
| ***N*** | 574 | 133 | 65 | 52 | 387 | 1,211 |
| ***Female (%)*** | 72 | 89 | 55 | 60 | 23 | 57 |
| ***Year of birth (median, IQ range)*** | 1943 (1934-1951) | 1961 (1945-1972) | 1951 (1940-1963) | 1950 (1940-1958) | 1935 (1928-1944) | 1942 (1932-1952) |
| ***Follow-up years (median, IQ range)**** | 15 (10-24) | 15 (10-19) | 17 (12-21) | 14 (10-19) | 18 (15-29) | 16 (12-25) |

**the follow-up years are the number of years between the first and the last ICD code of an individual.*

#### 3. Setting II: Patient selection through review of medical records

##### **Fig. S4** Flow patient selection in setting II

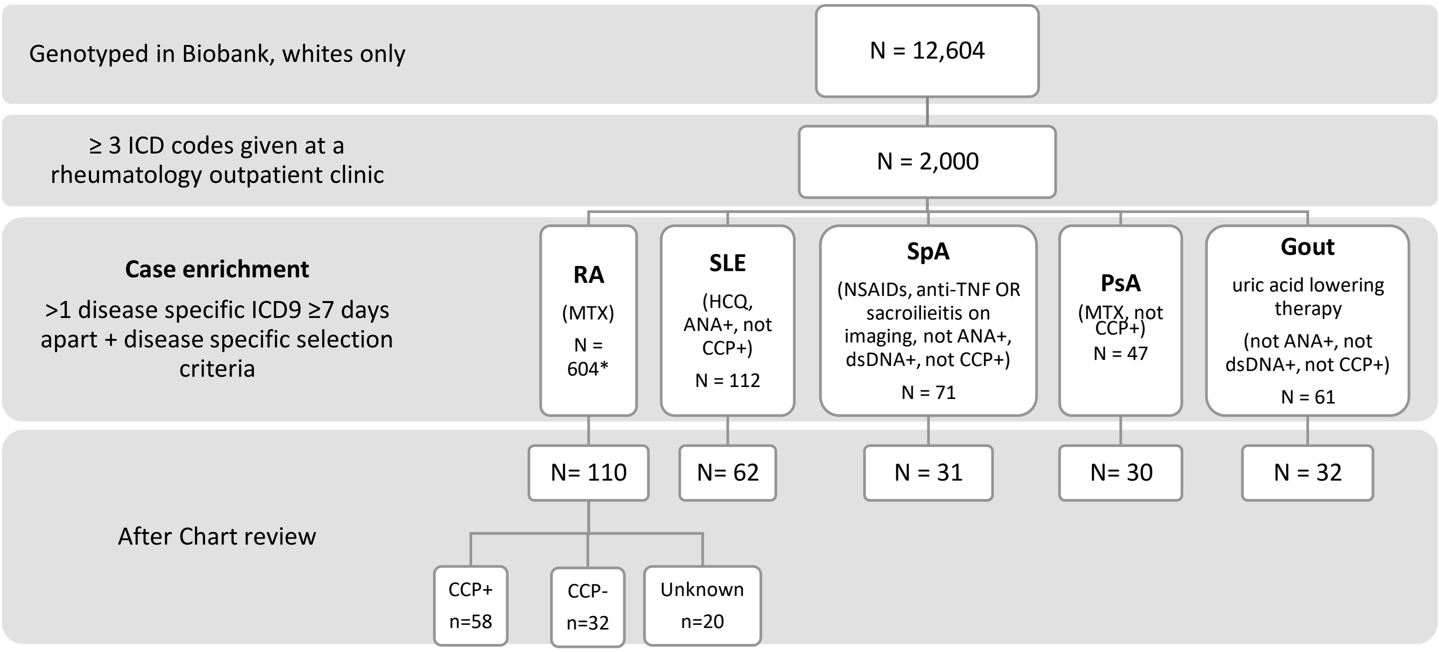

*HCQ = hydroxychloroquine; NSAIDs = non-steroid anti-inflammatory drugs; Anti-TNF = tumor necrosis factor inhibitors ;*

*CCP = cyclic citrullinated peptide antibody; ANA = antinuclear antibody; dsDNA= anti-double stranded DNA antibodies*

** not all patients were reviewed, because sufficient number of patients collected. Medical records were reviewed in random order*

**Table S4** Patient characteristics setting II Partners Biobank

| | Patients included in this study | | | | | | |
|---|---|---|---|---|---|---|---|
| | RA CCP+ | RA CCP- | SLE | SpA | PsA | Gout | Total patients |
| ***N*** | 58 | 32 | 62 | 31 | 30 | 32 | 245 |
| ***Female %*** | 88 | 78 | 89 | 26 | 63 | 25 | 68 |
| ***Year of Birth (median, IQ range)*** | 1953 (1944-1961) | 1951 (1945-1964) | 1962 (1953-1972) | 1964 (1956-1975) | 1950 (1946-1961) | 1942 (1936-1949) | 1955 (1945-1967) |
| ***Median follow-up duration notes (years, IQ range)**** | 8 (4-11) | 8 (5-11) | 12 (4-18) | 10 (3-15) | 11 (6-16) | 5 (4-9) | 8 (4-13) |

#### 4. Setting III: Prospective patient collection

**Fig. S5** Flow patient selection in setting III

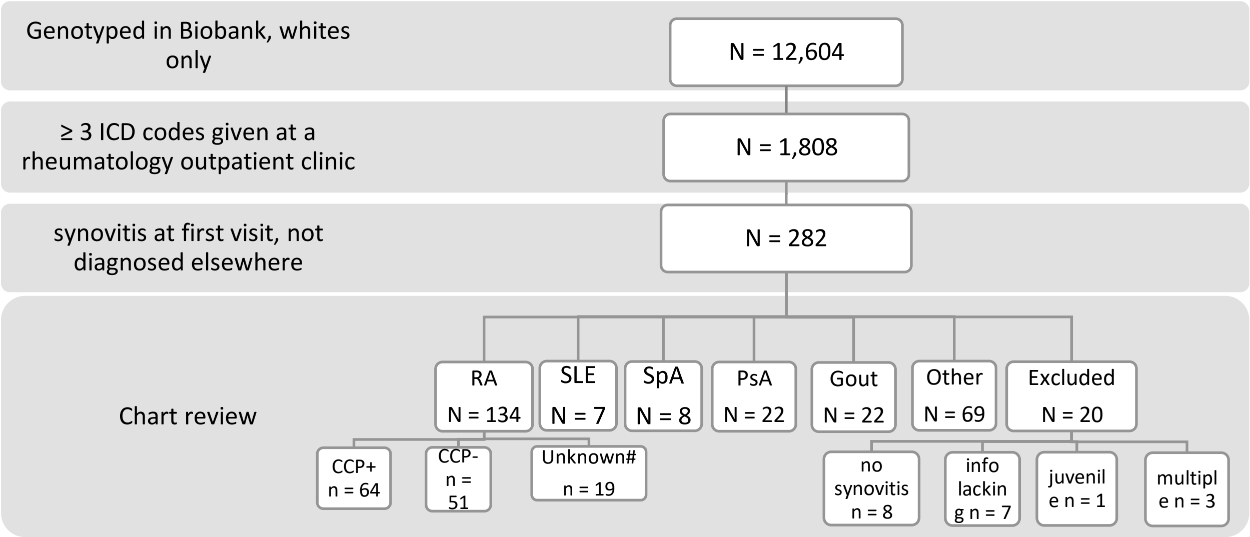

*RA = rheumatoid arthritis, SLE = systemic lupus erythematosus, SpA = spondyloarthropathy, PsA = psoriatic arthritis*

*# Unknowns were excluded from this analysis*

#### **Table S5** Patient characteristics setting III

| | Patients included in this study | | | | | | | |
|---|---|---|---|---|---|---|---|---|
| | RA CCP+ | RA CCP- | SLE | SpA | PsA | Gout | Other | Total within patients |
| **N** | 64 | 51 | 7 | 8 | 22 | 22 | 69 | 243 |
| **Female (%)** | 79 | 76 | 71 | 63 | 32 | 13 | 72 | 68 |
| **Year of birth (median, IQ range)** | 1953 (1945-1963) | 1950 (1943-1961) | 1962 (1961-1965) | 1976 (1961-1981) | 1965 (1948-1974) | 1944 (1941-1951) | 1952 (1942-1964) | 1953 (1943-1965) |
| **Follow-up duration (median yrs., IQ range)*** | 8 (4-11) | 8 (4-11) | 8 (3-14) | 6 (5-7) | 10 (4-13) | 3 (2-5) | 4 (2-8) | 7 (3-11) |

| | Excluded from the study | |
|---|---|---|
| | Excluded after medical record review | RA but no CCP info |
| **N** | 20 | 19 |
| **Female (%)** | 47% | 63% |
| **Year of birth (median, range)** | 1947 (1946-1954) | 1953 (1945-1958) |
| **Follow-up duration (median yrs., IQ range)*** | 11 (5-14) | 15 (11-17) |

** notes were extracted in 2017*

*RA = rheumatoid arthritis, SLE = systemic lupus erythematosus, SpA = spondyloarthropathy, PsA = psoriatic arthritis, Other = other rheumatic disease with synovitis*

**Fig. S6** Flow chart of the procedure of medical record review

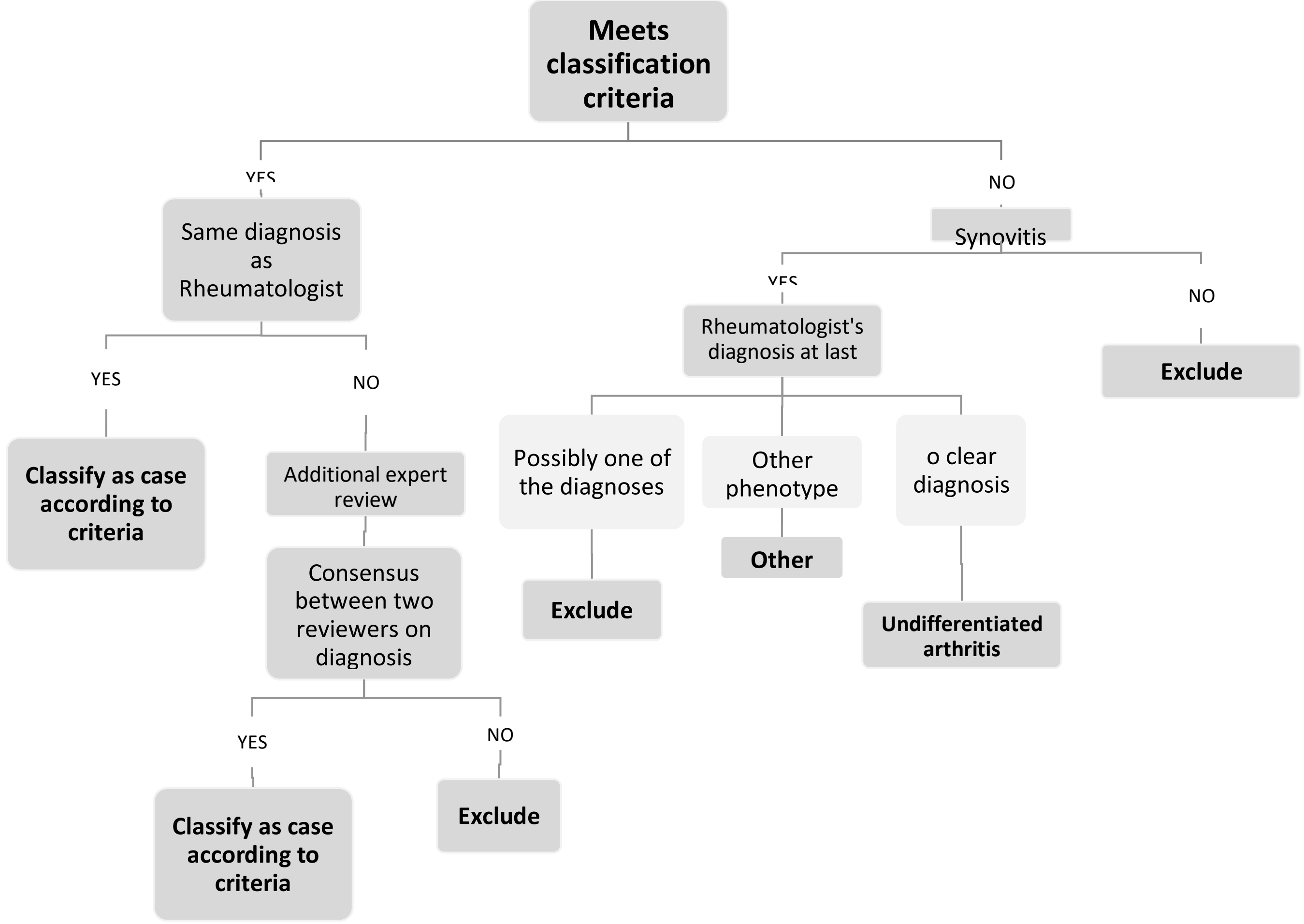

#Excluded patients because no clear decision could be made on whether the patient had undifferentiated arthritis or one of the diseases of our interest: either the rheumatologist diagnosed the patients without meeting the criteria (making it undifferentiated arthritis for our study) or the rheumatologist had more information than registered in the notes.

#### **Supplement 2** Sensitivity analyses

##### **Fig. S7** Sensitivity analysis A: performance of G-Prob on disease level

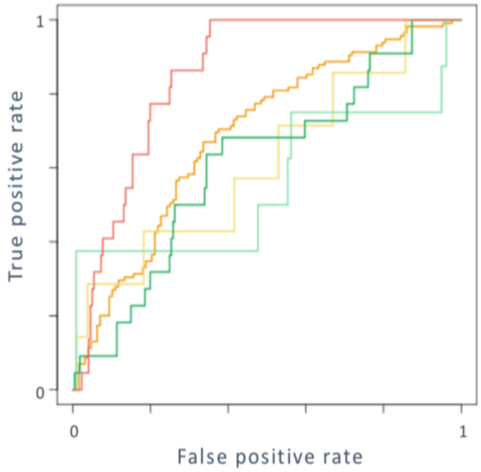

| | AUC | P value compared to complete model |
|---|---|---|
| RA | 0.75 (0.68-0.81) | *P* = 0.13 |
| SLE | 0.79 (0.72-0.85) | *P* = 0.60 |
| SpA | 0.87 (0.76-0.96) | *P* = 0.25 |
| PsA | 0.71 (0.63-0.82) | *P* = 0.07 |
| Gout | 0.82 (0.73-0.94) | *P* = 0.82 |

*RA = rheumatoid arthritis, SLE = systemic lupus erythematosus, SpA = spondyloarthropathy, PsA = psoriatic arthritis*

This graph depicts the receiver operating curve (ROC) from **Fig. 2B** (main manuscript) subdivided for each individual disease in setting II. The table shows area under the curve (AUC) for each disease.

##### **Fig. S8** Sensitivity analysis B: influence of individual diseases on G-Prob's performance.

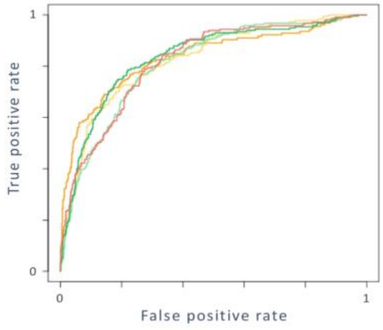

| | AUC | Performance compared to complete model |
|---|---|---|
| Excl RA | 0.83 (0.77-0.87) | *P* = 0.28 |
| Excl SLE | 0.82 (0.79-0.86) | *P* = 0.41 |
| Excl SpA | 0. 81 (0.78-0.84) | *P* = 0.81 |
| Excl PsA | 0.84 (0.80-0.87) | *P* = 0.20 |
| Excl Gout | 0.82 (0.81-0.82) | *P* = 0.56 |

*RA = rheumatoid arthritis, SLE = systemic lupus erythematosus, SpA = spondyloarthropathy, PsA = psoriatic arthritis*

This graph depicts the receiver operating curve of G-Prob when each time one disease is removed from G-Prob's calculation in setting II.

**Fig. S9** Sensitivity analysis C: Comparison of different shrinkage factors

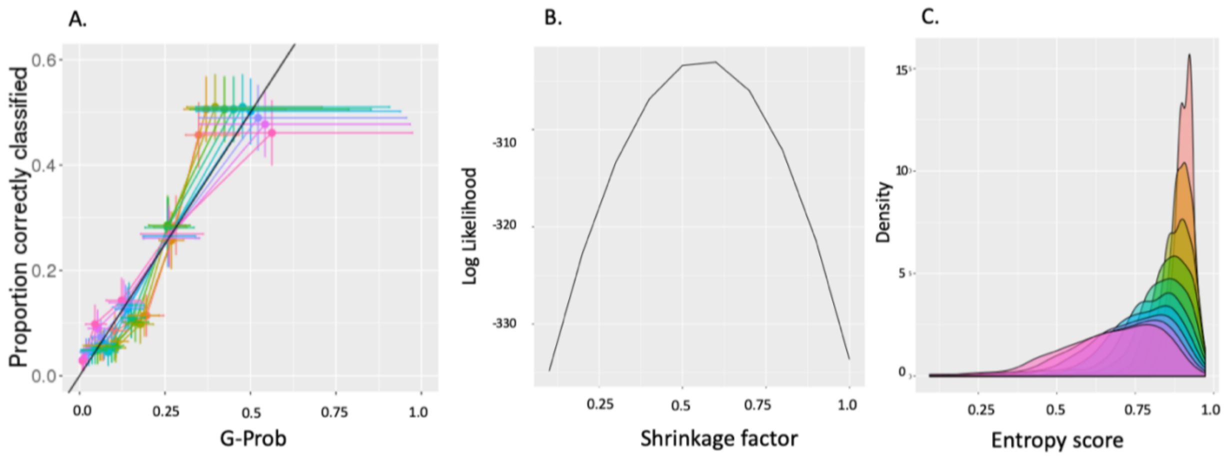

D.

| Shrinkage factor | Average entropy score | Proportion of patients with entropy ≤ 0.5 | 0.75 |
|---|---|---|---|
| 0.1 | 0.81 | 0 | 0 |
| 0.2 | 0.80 | 0 | 0.01 |
| 0.3 | 0.79 | 0 | 0.02 |
| 0.4 | 0.77 | 0.01 | 0.04 |
| 0.5 | 0.74 | 0.01 | 0.11 |
| 0.6 | 0.72 | 0.02 | 0.26 |
| 0.7 | 0.69 | 0.04 | 0.39 |
| 0.8 | 0.66 | 0.05 | 0.49 |
| 0.9 | 0.63 | 0.10 | 0.56 |
| 1 | 0.60 | 0.14 | 0.60 |

Plot A shows the results of setting II data with different shrinkage factors used to correct the logORs of the genetic risk scores. The x-axis displays the mean G-Prob (with range) of each quintile of G-Probs and the y-axis the corresponding proportion (with 95% confidence interval) of the G-Probs that concerned the patients' real disease. In the case of a perfect test performance, the lines would lie exactly on the black diagonal line.

Plot B gives the model fitness as expressed by the log likelihood of G-Prob with disease match for each different shrinkage factor. Here the higher the log likelihood the better the model fits.

Plot C shows the density of patients' entropy scores for probabilities created with different shrinkage factors.

Table D gives the average entropy score for G-Prob constructed with different shrinkage factors and the proportion of patients with an entropy score below 0.5 and 0.75.

**Supplement 3** Supplementary results

**Fig. S10** Density plots of G-Prob per disease

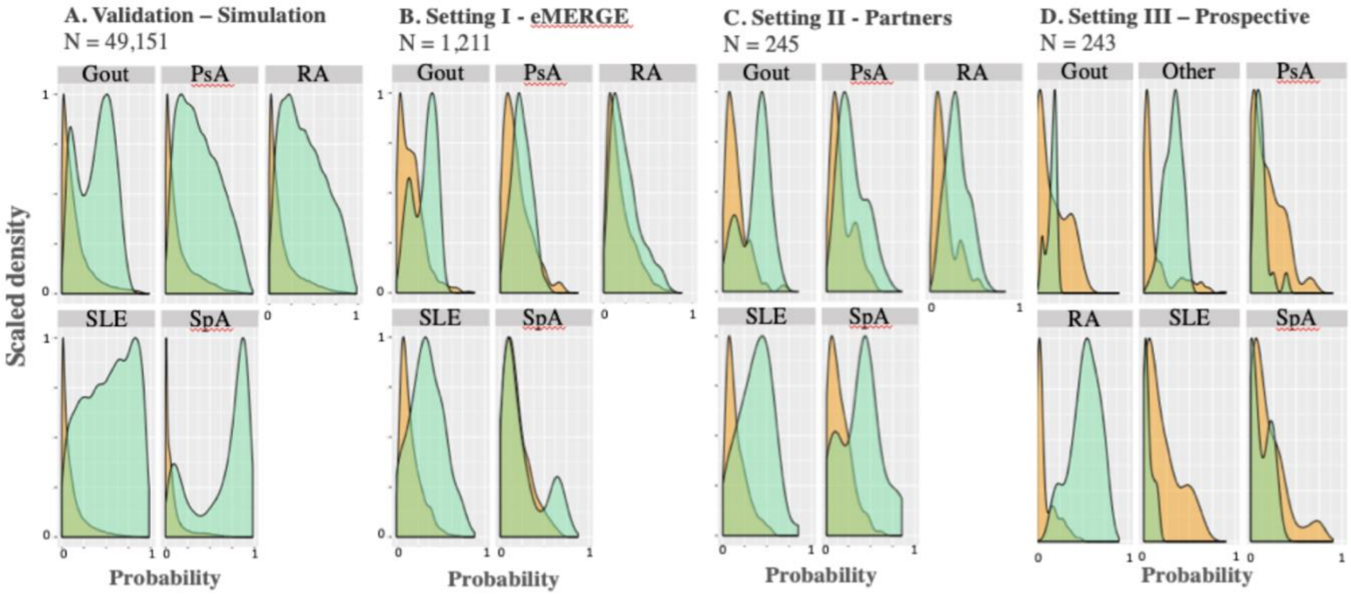

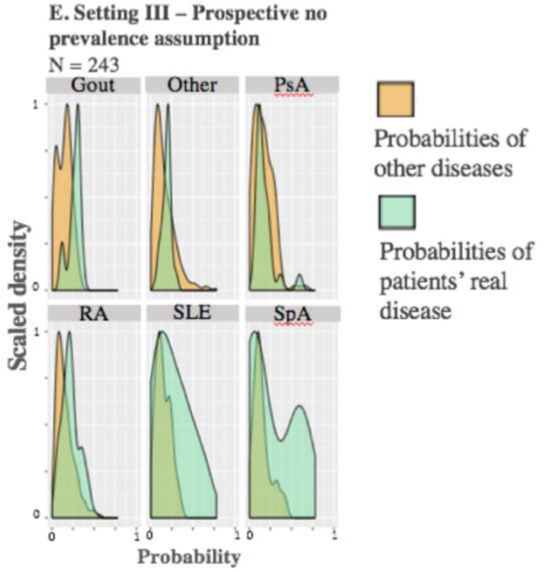

*RA = rheumatoid arthritis, SLE = systemic lupus erythematosus, SpA = spondyloarthropathy, PsA = psoriatic arthritis*

These graphs depict the density of probabilities for each disease subset in each setting (A-D). In green the probabilities that concern a patients' real disease. In orange the probabilities that refer to another disease than patients' real disease. Fig. E shows the results of the sub analysis of setting III where we applied a flat prevalence to G-Prob, avoiding skewed results due to an overrepresentation of (pre)RA cases.

**Table S6** Area under the receiver operating curve separated per disease

| | | ***AUC*** | ***95%CI*** | |
|---|---|---|---|---|
| ***Simulation*** | RA | 0.82 | 0.82 | 0.83 |
| | SLE | 0.90 | 0.90 | 0.91 |
| | SpA | 0.93 | 0.93 | 0.93 |
| | PsA | 0.81 | 0.81 | 0.82 |
| | Gout | 0.81 | 0.80 | 0.81 |
| | *macroAUC* | *0.86* | *0.85* | *0.86* |
| | *microAUC* | *0.86* | *0.86* | *0.86* |
| ***Setting I*** | RA | 0.69 | 0.65 | 0.72 |
| | SLE | 0.74 | 0.70 | 0.78 |
| | SpA | 0.58 | 0.50 | 0.67 |
| | PsA | 0.61 | 0.52 | 0.69 |
| | Gout | 0.78 | 0.75 | 0.80 |
| | *macroAUC* | *0.68* | *0.65* | *0.70* |
| | *microAUC* | *0.69* | *0.67* | *0.71* |
| ***Setting II*** | RA | 0.75 | 0.68 | 0.81 |
| | SLE | 0.79 | 0.72 | 0.85 |
| | SpA | 0.87 | 0.76 | 0.96 |
| | PsA | 0.71 | 0.63 | 0.82 |
| | Gout | 0.82 | 0.73 | 0.94 |
| | *macroAUC* | *0.79* | *0.74* | *0.82* |
| | *microAUC* | *0.81* | *0.76* | *0.84* |
| ***Setting III*** | RA | 0.69 | 0.63 | 0.76 |
| | SLE | 0.61 | 0.27 | 0.86 |
| | SpA | 0.56 | 0.33 | 0.84 |
| | PsA | 0.62 | 0.48 | 0.80 |
| | Gout | 0.85 | 0.80 | 0.91 |
| | Other | 0.57 | 0.51 | 0.66 |
| | *macroAUC* | *0.65* | *0.56* | *0.72* |
| | *microAUC* | *0.84* | *0.80* | *0.88* |

*microAUC = the AUC in the stacked dataset with 5 records per patient*

*macroAUC = the average of the AUC of all disease groups*

*RA = rheumatoid arthritis, SLE = systemic lupus erythematosus, SpA = spondyloarthropathy, PsA = psoriatic arthritis*

**Table S7** McFadden $R_2$ from multinomial logistic regression testing how much of the variance in the final diseases diagnosis is explained by clinical, genetic and serologic information.

| Independent variables | McFadden $R_2$ |
|---|---|
| genetic data | 17% |
| clinical data | 39% |
| serology | 31% |
| clinical + serology | 61% |
| clinical + genetic | 51% |
| genetic + clinic + serology | 73% |

Serologic testing is one of the first diagnostic steps a rheumatologist takes to differentiate between synovitis causing diseases. Though our research question focuses on the value of genetics before the first serology is ordered, we explored whether G-Prob would still improve the diagnostic accuracy when serologic information was available. So, we added the available CCP, RF, ANA and dsDNA serology information to the logistic regression analysis as factor coding for positivity, negativity and absence of the test.

A model with only clinical and serologic information had a $R_2$ of 61% which improved to 73% ($P<0.0001$) when our G-Prob was added (12% improvement). This demonstrates that G-Prob contributes independent information that exceeds the information obtained with initial examination and serologic testing.

**Figure S11** Test characteristics for the probabilities at different cut-offs.

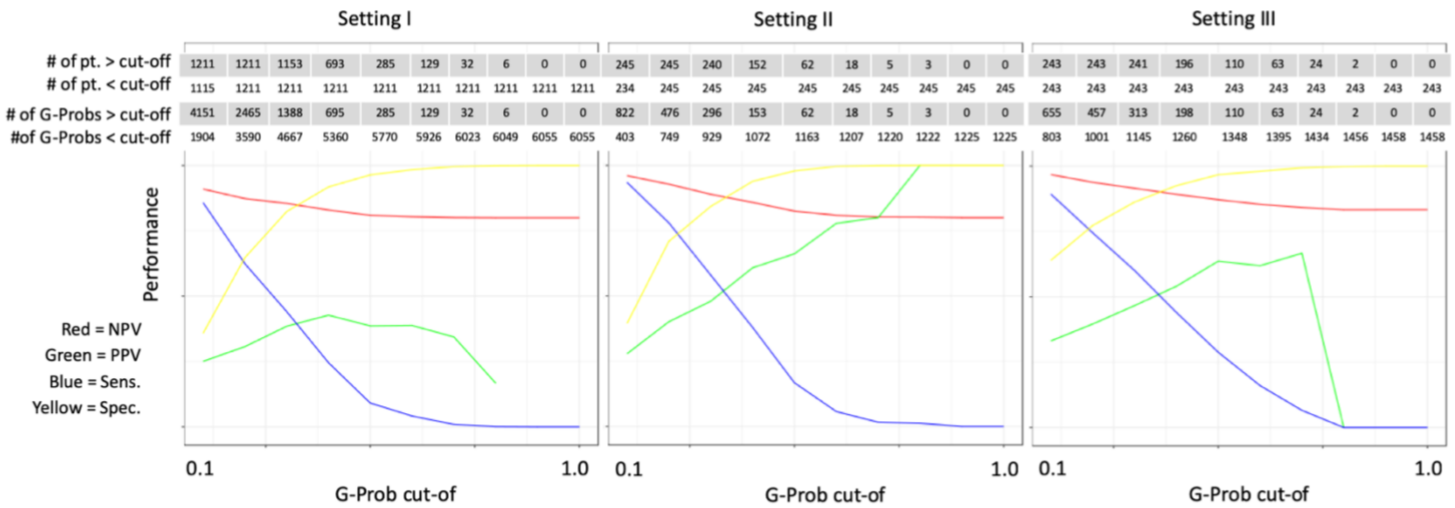

Setting I

| | | | | | | | | | | |
|---|---|---|---|---|---|---|---|---|---|---|
| # of pt. > cut-off | 1211 | 1211 | 1153 | 693 | 285 | 129 | 32 | 6 | 0 | 0 |
| # of pt. < cut-off | 1115 | 1211 | 1211 | 1211 | 1211 | 1211 | 1211 | 1211 | 1211 | 1211 |
| # of G-Probs > cut-off | 4151 | 2465 | 1388 | 695 | 285 | 129 | 32 | 6 | 0 | 0 |
| #of G-Probs < cut-off | 1904 | 3590 | 4667 | 5360 | 5770 | 5926 | 6023 | 6049 | 6055 | 6055 |

Setting II

| | | | | | | | | | | |
|---|---|---|---|---|---|---|---|---|---|---|
| # of pt. > cut-off | 245 | 245 | 240 | 152 | 62 | 18 | 5 | 3 | 0 | 0 |
| # of pt. < cut-off | 234 | 245 | 245 | 245 | 245 | 245 | 245 | 245 | 245 | 245 |
| # of G-Probs > cut-off | 822 | 476 | 296 | 153 | 62 | 18 | 5 | 3 | 0 | 0 |
| #of G-Probs < cut-off | 403 | 749 | 929 | 1072 | 1163 | 1207 | 1220 | 1222 | 1225 | 1225 |

Setting III

| | | | | | | | | | | |
|---|---|---|---|---|---|---|---|---|---|---|
| # of pt. > cut-off | 243 | 243 | 241 | 196 | 110 | 63 | 24 | 2 | 0 | 0 |
| # of pt. < cut-off | 243 | 243 | 243 | 243 | 243 | 243 | 243 | 243 | 243 | 243 |
| # of G-Probs > cut-off | 655 | 457 | 313 | 198 | 110 | 63 | 24 | 2 | 0 | 0 |
| #of G-Probs < cut-off | 803 | 1001 | 1145 | 1260 | 1348 | 1395 | 1434 | 1456 | 1458 | 1458 |

These figures depict the test characteristics (positive predictive value in green, negative predictive value in red, sensitivity in blue and specificity in yellow) of G-Prob's probabilities for different cut-offs (0.1 to 1.0 with increments of 0.1). The tables in each graph give the number of probabilities above and below the cut-off. Since each patient has multiple probabilities the tables also provide the number of patients that have probabilities above and below the cut-offs.

**Figure S12** Test characteristics for the probabilities at different cut-offs.

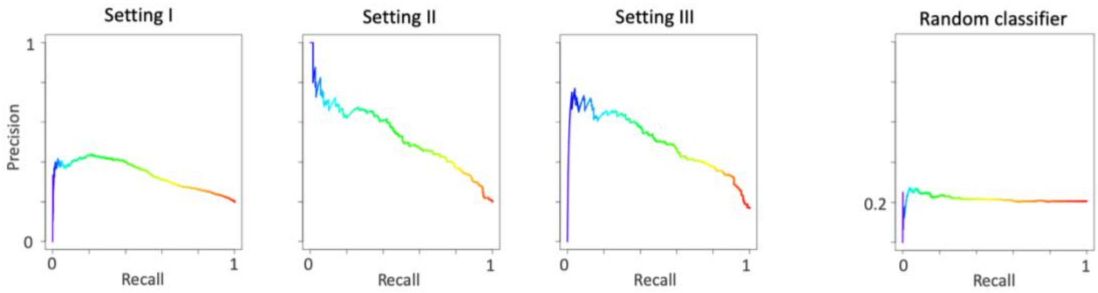

These graphs depict the PRC which is the precision (positive predictive value) versus recall (sensitivity) curve. The fourth graph is the PRC given a random classifier given a disease prevalence of 20% such as the case in the datasets of our study.
